# Human T cells employ conserved AU-rich elements to fine-tune IFN-γ production

**DOI:** 10.1101/830042

**Authors:** Julian J. Freen-van Heeren, Branka Popović, Aurélie Guislain, Monika C. Wolkers

## Abstract

Long-lasting CD8^+^ T cell responses are critical in combatting infections and tumors. The pro-inflammatory cytokine IFN-γ is a key effector molecule herein. We recently showed that in murine T cells, the production of IFN-γ is tightly regulated through AU-rich elements (AREs) that are located in the 3’ Untranslated Region (UTR). Loss of AREs resulted in prolonged cytokine production in activated T cells and boosted anti-tumoral T cell responses. Here, we investigated whether these findings can be translated to primary human T cells. Utilizing CRISPR-Cas9 technology, we deleted the ARE region from the *IFNG* 3’UTR in peripheral blood-derived human T cells. Loss of AREs stabilized the *IFNG* mRNA in T cells and supported a higher proportion of sustained IFN-γ protein-producing T cells. Importantly, this was also true for tumor antigen-specific T cells. MART-1 TCR engineered T cells that were gene-edited for ARE-deletion showed increased percentages of IFN-γ producing MART-1-specific ARE-Del T cells in response to MART-1 expressing tumor cells. Combined, our study reveals that ARE-mediated post-transcriptional regulation is highly conserved between murine and human T cells. Furthermore, generating antigen-specific ARE-Del T cells is feasible, a feature that could potentially be exploited for therapeutical purposes.

## INTRODUCTION

CD8^+^ T cells are critical for immunosurveillance and for the protection against invading pathogens. To do so, they produce effector molecules, including granzymes, chemokines, and cytokines. Interferon γ (IFN-γ) is a key cytokine for CD8^+^ T cells to exert their effector function [1]–[3]. IFN-γ is a pleiotropic cytokine that modulates angiogenesis, hematopoiesis, myelopoiesis and immune cell function [4]–[7]. During infections, IFN-γ can suppress the growth of pathogens through upregulation of antiviral factors [8], and attract myeloid cells such as neutrophils to the site of infection [9],[10]. Furthermore, IFN-γ sensing potentiates the innate immune response of dendritic cells, macrophages, monocytes and neutrophils to pathogens against (intra)cellular pathogens [9]–[13]. Indeed, point mutations and deletions in humans in the receptor for IFN-γ, *IFNGR1* and *IFNGR2*, which often lead to premature stop codons, revealed that IFN-γ sensing is instrumental to protect the host from infections by *Mycobacteria* species [14]–[16].

IFN-γ also prevents the development of cancers. In fact, mice lacking the *Ifng* gene, or the signaling protein downstream of Ifngr1/2, Stat1, spontaneously develop tumors [17],[18]. Furthermore, a high *IFNG*-mediated gene signature correlates with clinical response rates to immunotherapy in humans [19],[20]. Conversely, copy number alterations of IFN-γ pathway genes correlates with poor response to immunotherapy [21].

The regulation of IFN-γ production is multi-layered. The *IFNG* locus is only demethylated in effector and memory T cells [22], allowing for locus accessibility and transcription upon T cell activation. While the production of T cell effector molecules has been mainly attributed to changes in transcription and the presence of transcription factors [23]–[27], recently, the role of post-transcriptional regulation in T cells has also become appreciated [28]–[33]. Post-transcriptional regulation is mediated by sequence elements and structures present in both the 5’ and 3’ untranslated regions (UTRs) of mRNA molecules [34]–[37] and nucleoside modifications, such as adenine methylation [38]. By facilitating the binding of RNA binding proteins (RBPs), microRNAs and long non-coding RNAs, these regulators combined determine the actual protein output of a cell [37].

One of these sequence elements are adenylate uridylate-rich elements (AREs). AREs are AUUUA pentamers present in multimers in the 3’UTR of mRNA molecules [39],[40]. Interestingly, many cytokine transcripts contain AREs [37],[39]. They function as binding hubs for RBPs and microRNAs [39]–[41]. Binding to AREs by these factors mediate mRNA stability, localization and translation and thus fine-tunes the protein output [30],[41]–[44]. By deleting the 3’UTR AREs from cytokine mRNA, the protein production is decoupled from ARE-mediated post-transcriptional regulation [30],[43],[45]. We recently showed that AREs present in the *Ifng* 3’UTR dampen anti-tumoral responses in a murine melanoma model [46]. In fact, removal of AREs from the *Ifng* locus resulted in prolonged IFN-γ production in a tumor suppressive microenvironment, which translated into substantially delayed tumor outgrowth and prolonged survival [46].

The 3’UTR of IFN-γ is highly conserved between mice and men, in particular the region containing the AREs [30]. Therefore, we hypothesized that the regulation of IFN-γ production is also conserved. To unravel the post-transcriptional regulation of IFN-γ in primary human T cells, we removed a 160bp region by CRISPR-Cas9 technology from the *IFNG* 3’UTR locus that contained all AREs sequences. Similar to murine *Ifng* [46], removal of AREs from the human *IFNG* locus (ARE-Del) results in increased IFN-l production. Combining TCR gene transfer with ARE-deletion in primary human T cells confirmed increased IFN-γ production by ARE-Del T cells in response to tumor cells. The ARE-mediated regulation of IFN-γ is thus conserved in human T cells.

## MATERIALS AND METHODS

### Human PBMCs and cell culture

Studies with human T cells from anonymized healthy donors were performed in accordance with the Declaration of Helsinki (Seventh Revision, 2013) after written informed consent (Sanquin). Peripheral blood mononuclear cells (PBMCs) were isolated through Lymphoprep density gradient separation (Stemcell Technologies). Cells were used after cryopreservation.

Human T cells were cultured in T-cell mixed media (Miltenyi) supplemented with 5% heat-inactivated human serum (Sanquin) and fetal bovine serum (FBS, Bodinco), 2mM L-glutamine, 20 IU/mL penicillin G sodium salts, 20 μg/mL streptomycin sulfate (all Sigma Aldrich), 100 IU/mL recombinant human (rh) IL-2 (Proleukin, Novartis) and 10 ng/mL rhIL-15 (Peprotech), and were cultured in a humidified incubator at 37 °C + 5% CO_2_. Cells were cultured at a density between 0.5 – 1*10^6^ cells/mL. Medium was refreshed every 3 days.

### T cell activation

T cells were activated as previously described [47]. Briefly, 24-well plates were pre-coated overnight with 0.5 μg/mL rat α-mouse IgG2a (clone MW1483, Sanquin) at 4 °C. Plates were washed once with PBS, coated with 1 μg/mL α-CD3 (clone Hit3a, eBioscience) for a minimum of 2h at 37 °C, and washed once with PBS. 1*10^6^ PBMCs/well were added in medium supplemented with 1 μg/mL α-CD28 (clone CD28.2, eBioscience). Cells were activated for 72h in a humidified incubator at 37 °C and 5% CO_2_.

### crRNA and sequence primer design

crRNAs and sequence primers were designed using the CRISPR and Primer design tools in Benchling (https://benchling.com, ***Table 1***). Sequences were verified to be specific for the target of interest via BlastN and PrimerBlast (both NCBI).

**Table 1.**
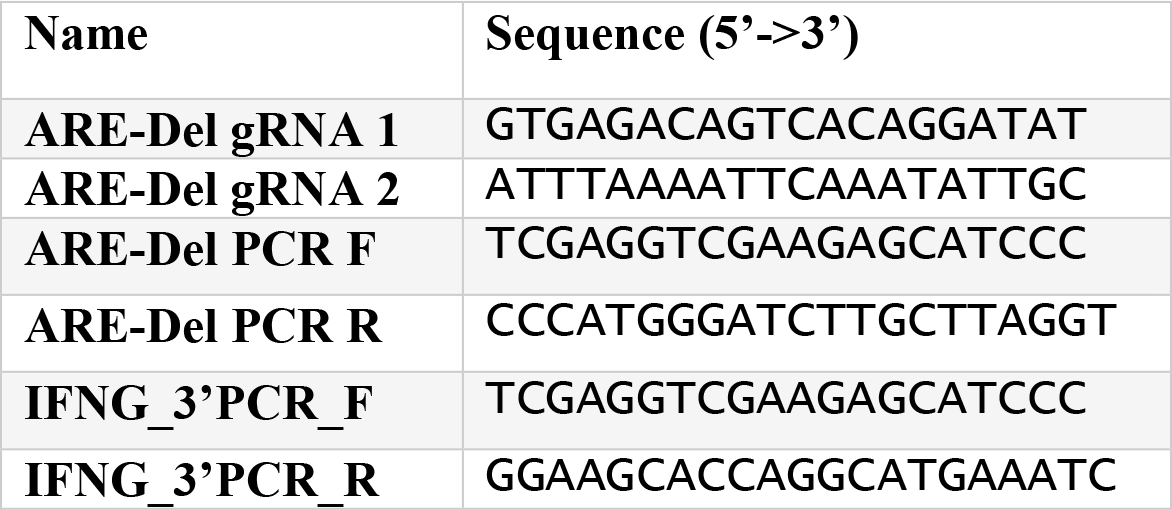
Primers used in this study.

### Genetic modification of human T cells with Cas9-RNPs

Cas9 RNP production and T cell nucleofection was performed as previously described [48]. Briefly, Alt-R crRNA and ATTO550-labeled tracr-RNA were reconstituted to 100 μM in Nuclease Free Duplex buffer (all Integrated DNA Technologies). As a negative control, non-targeting negative control crRNA #1 was used (Integrated DNA Technologies). Oligos were mixed at equimolar ratios (i.e. 4.5 μL total crRNA + 4.5 μL tracrRNA) in nuclease-free PCR-tubes and denatured by heating at 95 °C for 5min in a thermocycler. Nucleic acids were cooled down to room temperature prior to mixing them via pipetting with 30 μg TrueCut Cas9 V2 (Invitrogen) to produce Cas9 ribonuclear proteins (RNPs). Mixture was incubated at room temperature for at least 10min prior to nucleofection.

For nucleofection, 1*10^6^ T cells/condition were harvested 72h after α-CD3/α-CD28 activation and transferred to DNA Lo-binding Eppendorf tubes (Eppendorf). Cells were washed once with PBS and supernatant was completely removed. Cells were resuspended in 20 μL P2 Buffer (Lonza), Cas9 RNPs were added, and incubated for 2min. Cells were then electroporated in 16-well strips in a 4D Nucleofector X unit (Lonza) with program EH-100. 100 μL of pre-warmed complete medium was added and cells were allowed to recover for 5min in a humidified incubator at 37 °C and 5% CO_2_. Cells were transferred to 48-well plates containing 500 μL pre-warmed complete medium and cultured in a humidified incubator at 37 °C + 5% CO_2_.

CRISPR-mediated gene editing was tested on genomic DNA. Snapfrozen cell pellets (1*10^6^ cells) were incubated overnight at 56 °C while rotating at 850 RPM in 50 μL lysis buffer (50 mM Tris-HCl, 2.5 mM EDTA, 50 mM KCl and 0.45% Tergitol at pH 8.0) that was freshly supplemented with 1 μg/mL proteinase K (Roche). After deactivating proteinase K by incubating for 15min at 95 °C, lysed cells were centrifuged for 10min at 13.000 RPM (20.000g). Supernatant was transferred to a new tube. The *IFNG* genomic locus was amplified by PCR (30s at 98 °C, (10s at 98 °C, 30s at 65 °C, 30s at 72 °C) ×40, 7min at 72 °C, 2min at 15 °C) with DreamTaq HotStart Green Polymerase (ThermoScientific) and primerpair IFNG_PCR_F and IFNG_PCR_R (***Table 1***) and subsequently analyzed on 0.8% agarose gel. For Sanger sequencing, PCR product was purified with NucleoSpin Gel and PCR Clean-up kit (Macherey Nagel) according to manufacturer’s protocol (Baseclear).

ARE-deletion from *IFNG* mRNA was determined by total RNA extraction from snapfrozen pellets of 0.2*10^6^ cells activated for 3h with α-CD3/α-CD28 activated T cells using Trizol (Invitrogen). DNA contamination was removed with RNA Clean & Concentrator 5 kit (Zymo Research) according to manufacturer’s protocol. cDNA synthesis was performed in the presence or absence of SuperScript III Reverse Transcriptase according to manufacturer’s protocol (Invitrogen). The resulting cDNA was amplified by PCR (30s at 98 °C, (10s at 98 °C, 30s at 62 °C, 30s at 72 °C)×40, 7min at 72 °C, 2min at 15 °C) with with DreamTaq HotStart Green Polymerase and primerpair IFNG_3’PCR_F and IFNG_3’PCR_R (***Table 1***) and subsequently analyzed on a 2% agarose gel.

### Generation of MART-1 TCR expressing T cells

PBMCs were activated for 72h with α-CD3/α-CD28, harvested, and transduced with MART-1 TCR retrovirus as previously described [49]. Briefly, non-tissue culture treated 24-well plates were coated with 50 μg/mL Retronectin (Takara) overnight at 4 °C and washed once with PBS. Subsequently, 300 μL viral supernatant was added per well and was centrifuged for 1h at 4°C at 4000 rpm (2820g). 1*10^6^ freshly activated T cells were added per well, spun for 10min at 1000 rpm (180g), and cultured at 37 °C and 5 % CO_2_ in a humidified incubator. After 24h, cells were harvested and cultured in T25 flasks at a concentration of 0.8*10^6^ cells/mL for 3 days in complete medium.

### Functional assays

T cells were stimulated with 1 μg/mL pre-coated α-CD3 and 1 μg/mL soluble α-CD28 for indicated time points. MART-1 TCR-transduced T cells were co-cultured with HLA-A*0201^+^ MART1^hi^ Mel 526 (MART1^+^), or HLA-A*0201^−^ MART1^lo^ Mel 888 (MART1^−^) tumor cell lines [50]–[52], in a 1:1 effector to target (E:T) ratio for indicated time points. 1 μg/mL Brefeldin A (BD Biosciences) was added as indicated. Non-activated T cells were used as a negative control. All stimulations were performed in T cell mixed medium supplemented with 10% FBS, 2mM L-glutamine, 20U/mL penicillin G sodium salts and 20μg/mL streptomycin sulfate.

### Flow cytometry and intracellular cytokine staining

For flow cytometric analysis, cells were washed with FACS buffer (PBS, containing 1% FBS and 2mM EDTA) and labeled with monoclonal antibodies α-CD4 (clone SK3), α-CD8 (clone SK1), α-murine TCR beta (clone H57-597, all BD Biosciences), α-IFN-γ (clone 4S.B3), αIL-2 (clone MQ1-17H12) and α-IFN-α (clone MAb11) (all Biolegend). Near-IR (Life Technologies) was used to exclude dead cells from analysis. For intracellular cytokine staining, cells were cultured in the presence of 1 μg/mL Brefeldin A for indicated timepoints and were fixed and permeabilized with Cytofix/Cytoperm kit (BD Biosciences) according to manufacturer’s protocol. Expression levels were acquired using FACSymphony (BD Biosciences) and data were analyzed using FlowJo (FlowJo LLC, version 10). For gating strategy, see ***Suppl Fig 1A-B***.

### Real-time PCR analysis

Total RNA was extracted using Trizol (Invitrogen). cDNA synthesis was performed with SuperScript III Reverse Transcriptase (Invitrogen), and real-time PCR was performed with SYBR green and a StepOne Plus RT-PCR system (both Applied Biosystems). Reactions were performed in duplicate or triplicate, and cycle threshold values were normalized to *18S* levels. Primer sequences for *IFNG, TNF, and IL2* mRNA were previously described [47].

To determine the half-life of *IFNG* mRNA, T cells were activated for 3 h with indicated stimuli, and then treated with 5 μg/ml actinomycin D (ActD) (Sigma-Aldrich). Data were analyzed using StepOne Plus software (Applied Biosystems).

### Statistical analysis

Results are expressed as mean ± SD when indicated. Statistical analysis between groups was performed with Prism (GraphPad Software, version 8), using paired or ratio paired Student’s t-test when comparing 2 groups, or two-way ANOVA with Bonferroni’s correction for multiple comparisons when comparing more than 2 groups. P values <0.05 were considered to be statistically significant.

## RESULTS

### Deletion of AREs from the *IFNG* locus by CRISPR-Cas9

The human *IFNG* 3’UTR contains 5 AU-rich elements (AREs), defined as AUUUA (***Figure 1A***, underlined sequence). To remove all 5 ARE sequences within the 3’UTR of the *IFNG* locus, we designed 2 CRISPR guide RNAs (crRNAs) (***Figure 1A***, bold sequence). As a control, we included non-targeting crRNAs (control). PBMC-derived human primary T cells were activated with α-CD3/α-CD28 for 3 days prior to nucleofection with 30 μg Cas9-RNP complexes (***Figure 1B***). Using ATTO-labeled tracrRNA allowed us to follow nucleofection efficiency by flow cytometry. 3 days after nucleofection, virtually all CD8^+^ and CD4^+^ T cells were positive for ATTO550 (control 97,8± 0,9 %, ARE-Del 98,6 ± 0,4 %, ***Figure 1C, D***). In line with the described cleavage pattern of *S. pyogenes* Cas9 [53], cleavage occurred between the 17^th^ and 18^th^ base of the designed crRNAs (***Suppl Figure 2A***). The 160 bp deletion containing the AREs was efficient, as revealed by both PCR and by Sanger sequencing of the genomic *IFNG* 3’UTR region (***Figure 1E***; ***Suppl Figure 2A***). Importantly, nucleofection did not impact T cell survival (***Figure 1D***), and ARE-Del nucleofected T cells expanded as efficiently as control nucleofected T cells, with a 9.2 ± 1.1 fold vs 8.9 ± 1.2 fold expansion within 8 days after removal from α-CD3/α-CD28.

**Figure 1.**
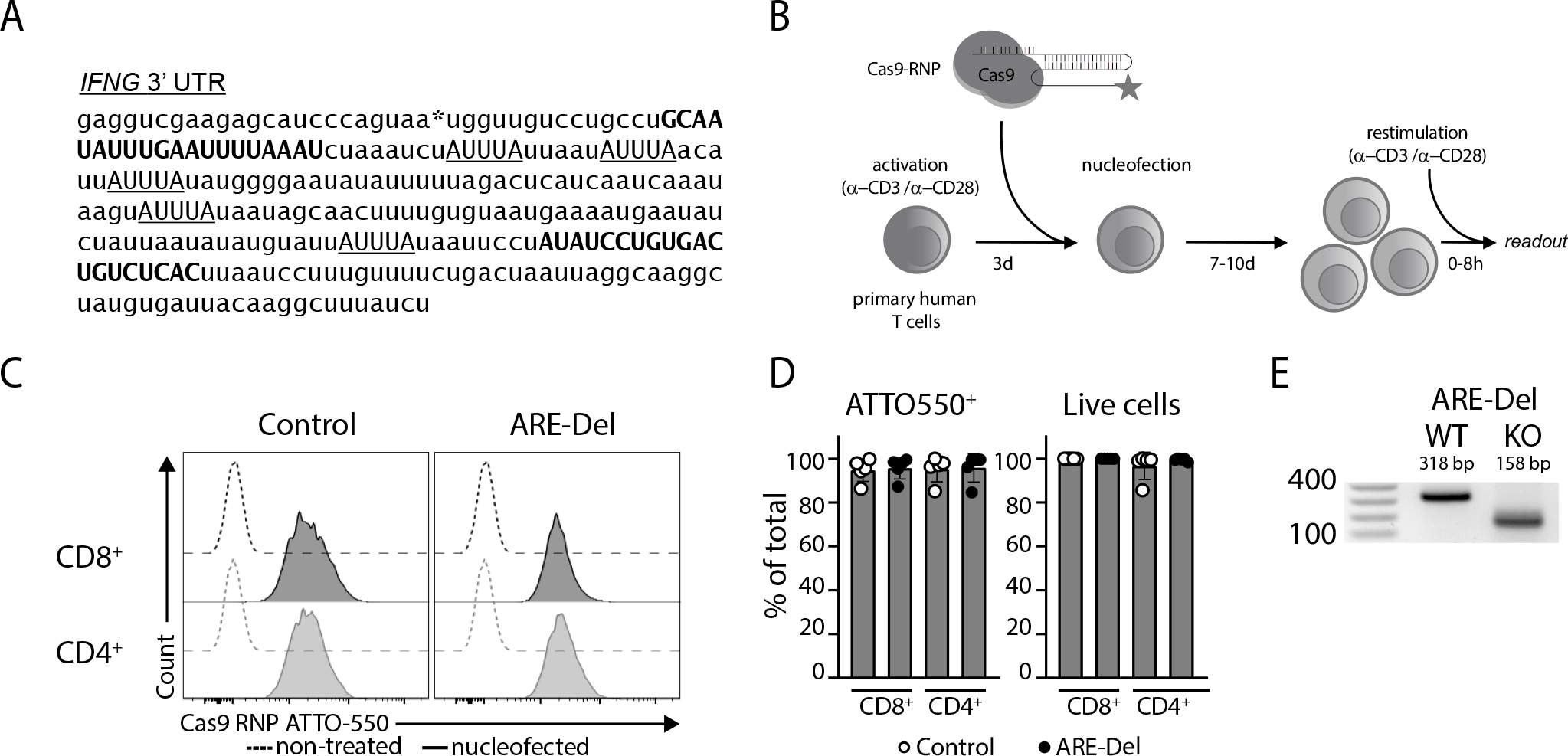
Generation of primary human ARE-Del T cells. **(A**) Sequence of the human *IFNG* 3’ UTR. Translation stop site is indicated with an asterisk, crRNA sequences are bold capitals, AREs are underlined capitals. **(B)** Graphical representation of the experimental setup. (**C** and **D**) PBMCs were stimulated for 72h with α-CD3/α-CD28 and subsequently nucleofected with Cas9 RNPs with ATTO550 labeled crRNAs targeting the 3’UTR of *IFNG* (ARE-Del), or with non-targeting control crRNA (Control). (C) Representative histograms of Cas9 RNP-ATTO 550 fluorescence gated on live CD8^+^ T cells (black) and live CD4^+^ T cells (gray). Dotted histograms represent non-nucleofected T cells. (D) Compiled data on ATTO550 fluorescence (left) and percentage of live cells (right) from n = 5 healthy donors from 3 independent experiments gated on CD8^+^ and CD4^+^ T cells, respectively. (**E**) ARE-deletion was analyzed by genomic PCR and subsequent gel analysis.

To determine if gene editing affected the *IFNG* 3’UTR downstream of the ARE region, we extracted RNA from control and ARE-Del T cells. T cells were first activated for 3 h with α-CD3/α-CD28 to increase *IFNG* mRNA levels. After cDNA synthesis, we performed PCR from the *IFNG* mRNA coding region to the 3’end of the *IFNG* 3’UTR, resulting in a 509 bp fragment for the WT and a 349 bp fragment for the ARE-Del *IFNG mRNA* (***Suppl Figure 2B***). The gene-editing was thus specific for the ARE region and did not result in a premature transcriptional stop of the *IFNG* 3’UTR. In conclusion, ARE-deletion from the *IFNG* locus was efficient and specific in primary human T cells and did not impact T cell survival and expansion.

### ARE-deletion stabilizes *IFNG* mRNA and augments protein production in human T cells

We next studied whether ARE-deletion from the *IFNG* locus altered *IFNG* mRNA levels. In T cells that were rested for 9 days after nucleofection, *IFNG* mRNA levels were approximately 2-fold higher in ARE-Del T cells compared to control T cells (***Figure 2A***). In most donors, *IFNG* mRNA levels of ARE-Del T cells were also slightly increased upon 3h stimulation with α-CD3/α-CD28 compared to control T cells (***Figure 2A***). *TNFA* and *IL2* mRNA levels remained unaltered, indicating *IFNG* mRNA specific alterations in ARE-Del T cells (***Suppl Figure 3A***). To determine *IFNG* mRNA half-life, T cells were treated with Actinomycin D, a drug that blocks *de novo* mRNA transcription. ARE-deletion significantly increased the stability of *IFNG* mRNA compared to control T cells (t1/2>2h compared to ~70min, ***Figure 2B***). This is similar to murine ARE-Del T cells [46].

**Figure 2.**
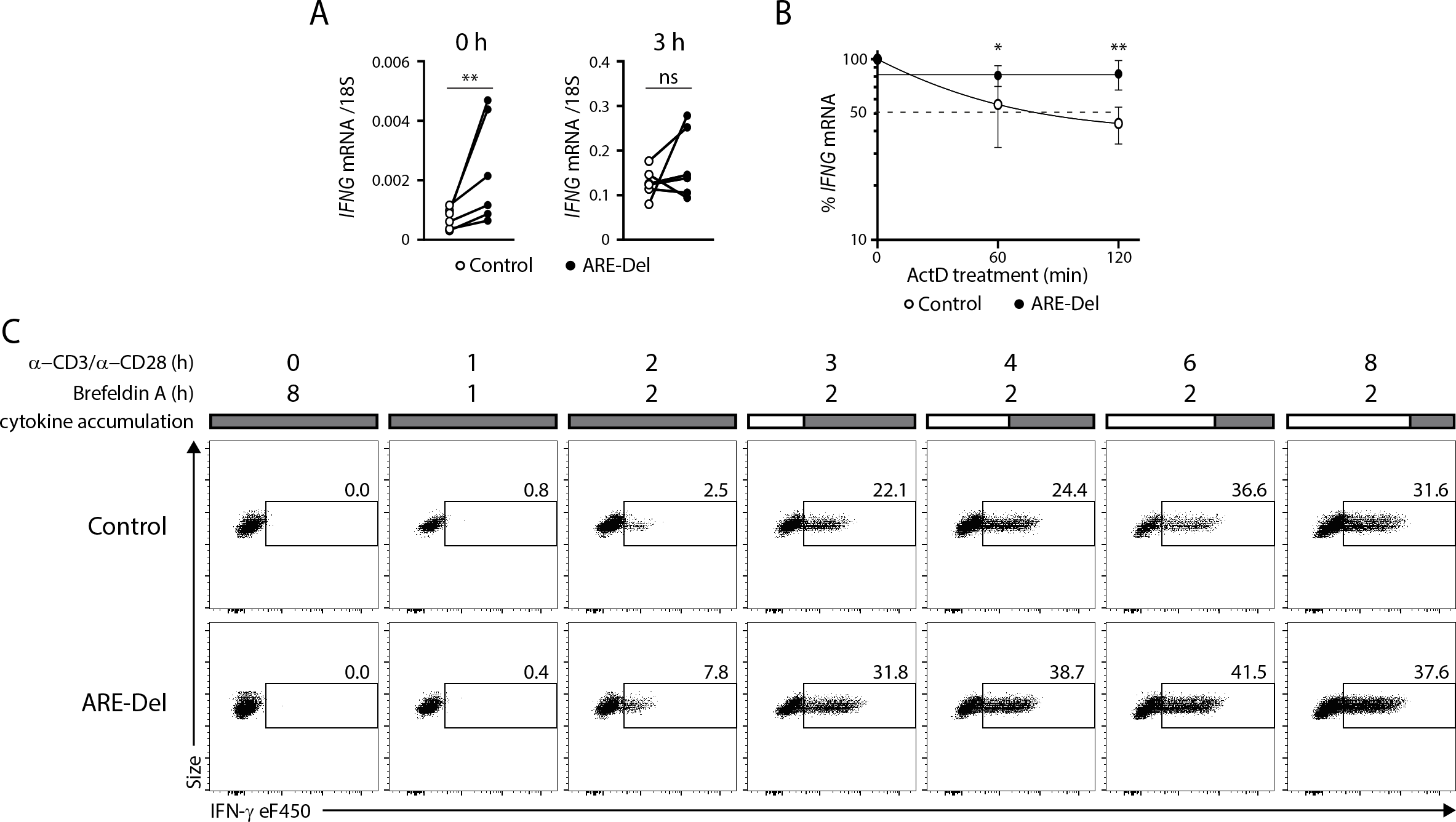
Enhanced *IFNG* mRNA levels, mRNA stability and protein output in ARE-Del T cells. (**A-B**) T cells rested 9 days and were left untreated or restimulated for 3h with α-CD3/α-CD28. (A) *IFNG* mRNA levels as determined by RT-PCR from n = 6 donors from 2 independent experiments (paired Student’s t-test; *p < 0.05, ns, not significant). (B) T cells were activated for 3h with α-CD3/α-CD28. Actinomycin D was added to block *de novo* mRNA transcription, and *IFNG* mRNA stability was determined following 0, 60 and 120 minutes of Actinomycin D treatment. n = 4 donors from 2 independent experiments (Two-way ANOVA; *p < 0.05, **p < 0.01). (**C** and **D**) Resting T cells were restimulated with α-CD3/α-CD28 in the presence of Brefeldin A for indicated time points. (C) Representative dot plots of IFN-γ production gated on live CD8^+^ T cells. (D) Compiled data on IFN-γ^+^ cells (top row) and IFN-c geoMFI (bottom row) gated on IFN-γ ^+^ CD8^+^ and CD4^+^ T cells from n = 6-13 donors from 5-7 independent experiments after 2, 4 and 6h of stimulation (paired Student’s t-test; *p < 0.05, **p < 0.01, ns, not significant). Brefeldin A was added during the last 2h of activation. T cells cultured in the absence of stimuli were used as negative control.

To determine whether ARE-deletion also modulated the IFN-γ protein production and/or kinetics, we activated ARE-Del and control T cells with α-CD3/α-CD28. To visualize the production kinetics, we added Brefeldin A for a maximum of the final 2h of stimulation [54], and measured the production of IFN-γ protein via intracellular cytokine staining. In non-activated T cells, we did not observe any protein production by any T cell type (***Figure 2C***), indicating that like control T cells, ARE-Del T cells require T cell activation for cytokine production. Irrespective of the ARE-deletion, the production of IFN-γ by T cells peaks at 6h after stimulation (***Figure 2C***), as previously described [3],[47]. However, already from 2h of activation onwards, we found increased percentages of IFN-γ-producing T cells in ARE-Del CD8^+^ T cells compared to control T cells (***Figure 2C, D***). This increase in cytokine production was specific to IFN-γ and was not observed for TNF-α or IL-2 (***Suppl Figure 3B, C***). Even though we did not find overt differences in the IFN-γ production per cell (***Figure 2D***, bottom row), deletion of AREs promoted the IFN-γ production in both CD8^+^ and in CD4^+^ T cells (***Figure 2D***, top row). Thus, ARE-deletion in the *IFNG* 3’UTR increases the stability of *IFNG* mRNA and augments the production of IFN-γ protein.

### Prolonged cytokine production by human ARE-Del T cells upon removal of stimuli

We next questioned if ARE-deletion also modulates mRNA expression and protein production kinetics upon removal of stimulation. We therefore restimulated ARE-Del T cells and control T cells with α-CD3/α-CD28 for 72h, and transferred the cells to new wells to remove the stimulus. 24h after removal of the stimulus, WT T cells expressed slightly lower *IFNG* mRNA levels than ARE-Del T cells, which further dropped after 48h (***Figure 3A***). In contrast, ARE-Del T cells maintained equally high *IFNG* mRNA levels for up to 72h (***Figure 3A***).

**Figure 3.**
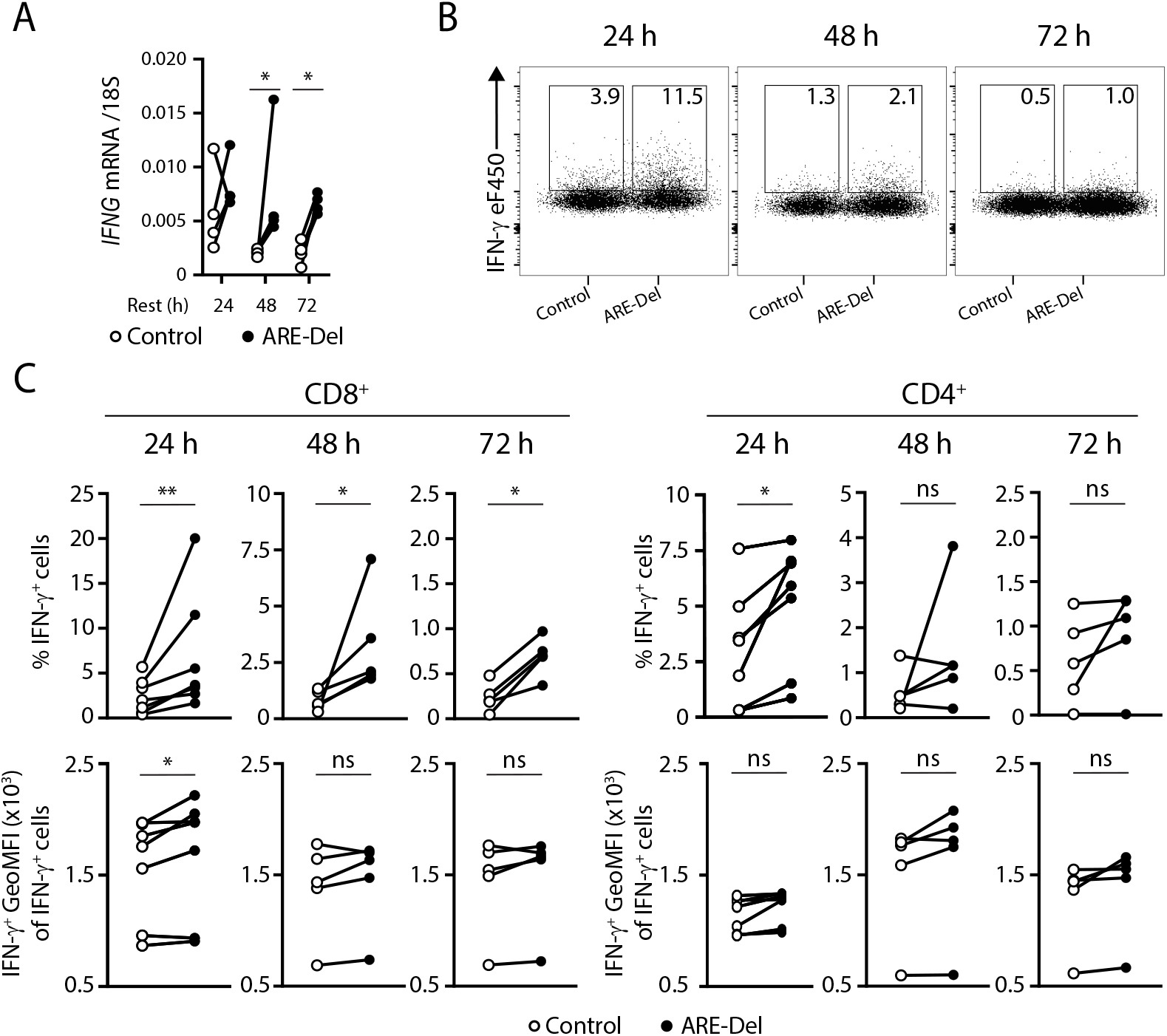
Prolonged IFN-γ production in ARE-Del T cells in the absence of stimulation. Rested T cells (11 days) were restimulated for 72h with α-CD3/α-CD28 and removed from stimulus for indicated time. (**A**) *IFNG* mRNA levels as determined by RT-PCR from n = 4 donors from 1 experiment (paired Student’s t-test; *p < 0.05, ns, not significant). (**B** and **C**) IFN-γ protein production was assessed by adding Brefeldin A 4h prior to IFN-γs production assessment. (B) Representative concatenated dot plots of IFN-γ production gated on live CD8^+^ T cells after removal from stimulus for indicated time points. (C) Compiled data on IFN-γ protein production from n = 7 donors from 3 independent experiments (Ratio paired Student’s t-test; *p < 0.05, ** p < 0.01, ns, not significant).

Intriguingly, also the IFN-γ protein production was altered by ARE-deletion. Even though TNF-α and IL-2 production was lost at 24h after removal from stimulation (***Suppl Figure 3D***), ARE-Del CD8^+^ T cells retained their capacity to produce IFN-γ for up to 72h (***Figure 3B, C***). This also translated in higher IFN-γ production per cell at 24h post antigen removal, as determined by the geoMFI of the IFN-γ producing T cells (***Figure 3C***). This increased production is also observed in CD4^+^ ARE-Del T cells, albeit only for the first 24h (***Figure 3C***). Together, these data show that, similar to murine T cells [30], deletion of AREs from the *IFNG* 3’ UTR results in prolonged IFN-γ production in human T cells.

### TCR-engineered ARE-Del T cells are superior IFN-γ producers in response to tumor cells

IFN-γ mediated signaling is key for effective anti-tumoral T cell responses [55]. This prompted us to study whether ARE-Del T cells are also superior IFN-γ producers when encountering tumor cells. To this end, we combined ARE-deletion with TCR gene transfer. We first retrovirally transduced human CD8^+^ T cells with a codon-optimized MART-1 TCR that recognizes the HLA-A*0201 restricted epitope of MART-1 (aa26-35) [49]. After 5 days, MART-1 TCR engineered T cells were restimulated with α-CD3/α-CD28 for 72h prior to nucleofection. Of note, ARE-deletion was also highly efficient in TCR-engineered T cells (***Suppl Figure 3E***). Furthermore, the combined TCR engineering with subsequent nucleofection did not affect the viability or the expansion capacity of ARE-Del versus control MART-1 TCR engineered T cells (92.3±2.3 vs 91.3±2.6 % live cells, and 12.9±5.2 vs 13.2±4.2 fold expansion within 7 days, respectively).

To study the production of IFN-γ in response to tumor cells, we exposed MART-1 TCR-engineered ARE-Del T cells or control T cells to a MART-1^hi^ HLA-A*0201^+^ melanoma cell line (MART-1^+^) and to a MART-1^lo^ HLA-A*0201^−^ melanoma cell line (MART-1^−^) [50]–[52]. As expected, MART-1 TCR-engineered T cells only produce cytokines when co-cultured with MART-1^+^ tumor cells (***Figure 4A***). Interestingly, after 6h of co-culture with MART-1^+^ cells, a significantly higher percentage of MART-1 TCR-engineered ARE-Del CD8^+^ T cells produced IFN-γ than the control MART-1 TCR-engineered T cells (***Figure 4A, B***). Again, the higher percentage of IFN-γ) producing cells did not translate into significantly increased cytokine production per cell, as determined by the GeoMFI of the IFN-γ^+^ T cells (***Figure 4B***).

**Figure 4.**
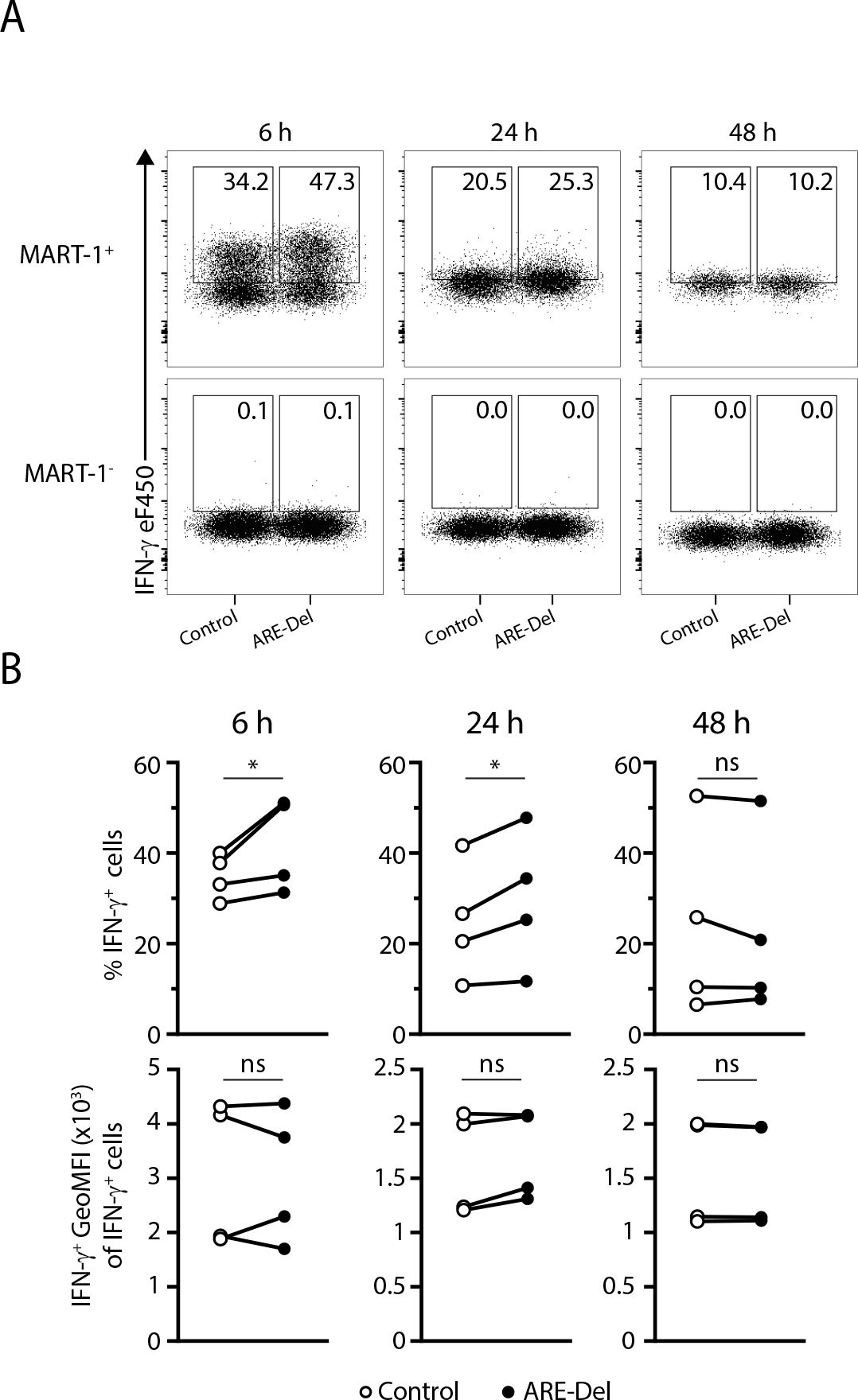
Enhanced IFN-γ production by ARE-Del T cells upon target cell recognition. MART-1 TCR-engineered T cells were cocultured with MART-1^+^ and MART-1^−^ cell lines expressing MART-1 peptide for indicated timepoints. Brefeldin A was added 2h prior to IFN-c production assessment. (**A**) Representative concatenated dot plots of IFN-) production gated on live CD8^+^ MART-1 TCR^+^ T cells after 6, 24 and 48h of co-culture. (**B**) Compiled data from n = 4 donors from 3 independent experiments (paired Student’s t-test; *p < 0.05, ns, not significant).

The increased percentage of IFN-γ producing cells in ARE-Del MART-1 TCR-engineered T cells compared to control T cells was also observed after 24h of co-culture (***Figure 4B***). T cells were retransferred to freshly seeded tumor cells for another 24h and cytokine production was measured. At this time point, i.e. 48h of co-culture, no differences in cytokine production were observed (***Figure 4B***). Overall, these finding show that the generation of antigen-specific ARE-Del T cells is feasible and allows to drive enhanced and prolonged IFN-γ production in response to target cells.

## DISCUSSION

Here we show that the deletion of AREs from the *IFNG* 3’UTR in primary human T cells results in higher *IFNG* mRNA levels, enhanced numbers of IFN-γ producing T cells and prolonged IFN-γ production. These findings are in line with previous findings with murine ARE-Del T cells [43],[46]. Thus, the highly conserved ARE region in the *IFNG* 3’UTR locus also provides conserved post-transcriptional regulation in T cells.

We also observed differences between murine and human ARE-Del T cells, which are primarily of quantitative nature. Murine ARE-Del T cells produce more IFN-γ per cell than wildtype T cells [46]. We did not observe this difference in human T cells. The overall mRNA stability in activated human control T cells is also lower than that of murine T cells. Whereas murine T cells show a t1/2>2h upon antigen stimulation [30],[33], human control T cells activated with α-CD3/α-CD28 only reach a t1/2 ~ 70min. This also translates into lower overall IFN-γ protein output in human T cells, both in terms of responding T cells (60-80% in murine wild type T cells versus 30-50% in human T cells upon PMA-Ionomycin stimulation [47],[54]), and in terms of production kinetics. In fact, even though human and murine T cells initiate the IFN-γ production in the same time frame, human T cells cease to produce IFN-γ faster. To date, it remains unresolved whether these differences stem from different activation methods in murine and human T cells (MEF cells expressing antigen and B7.1 in murine [46] versus α-CD3/α-CD28 in human T cells), from different T cell responsiveness due to different requirements on nutrients and stimuli during culture conditions, from higher diversity in human CD8^+^ T cell populations [56], or from intrinsic differences between murine and human T cells. Irrespective of the observed quantitative differences, we show here that ARE sequences are essential for the regulation of *IFNG* mRNA stability and IFN-γ protein output in both murine [46] and human T cells.

Excessive production of IFN-γ can induce immunopathology [57]–[65]. However, the AREs solely fine-tune post-transcriptional regulation of IFN-γ production, while leaving the requirement of antigen recognition and transcriptional regulation in murine and human T cells intact ([46], this study). Indeed, in studies with long term memory ARE-Del T cells in a *Listeria* infection and in a B16 melanoma model, we find the increased and prolonged IFN-γ production also only in the presence of tumor cells or after antigenic challenge [46]. We therefore anticipate only limited side effects of the ARE-deletion in T cells.

We here combined retroviral TCR engineering with efficient CRISPR-Cas9 mediated genome editing. This experimental setup is a powerful tool to study gene modification in human T cells in an antigen-specific setting. Furthermore, TCR engineering with CRISPR-Cas9 technology could also be useful for therapeutic purposes. It is noteworthy that upon CRISPR-Cas9 mediated genome editing, TCR-engineered T cells maintained their capacity to expand, a feature that is critical for generating large numbers of T cells required for cellular products.

In conclusion, the generation of primary human ARE-Del T cells revealed that fundamental post-transcriptional mechanisms of *IFNG* regulation are conserved throughout evolution.

## AUTHOR CONTRIBUTIONS / ACKNOWLEDGEMENTS

JJFH, BP, and MCW designed experiments, JJFH, BP and AG performed experiments, JJFH, BP and MCW analyzed data, and JJFH and MCW wrote the manuscript. MCW supervised the project.

The authors would like to thank Dr. Ton Schumacher (Netherlands Cancer Institute) for providing MART-1 TCR viral supernatants, Nordin Zandhuis for technical assistance, and the Wolkers’ lab for scientific input and critical reading of the manuscript.

This research was supported by KWF Kankerbestrijding (10132), the Dutch Science Foundation (VIDI grant 917.14.214), and Oncode Institute, all to M.C.W.

---

**Supplemental Figure 1.**
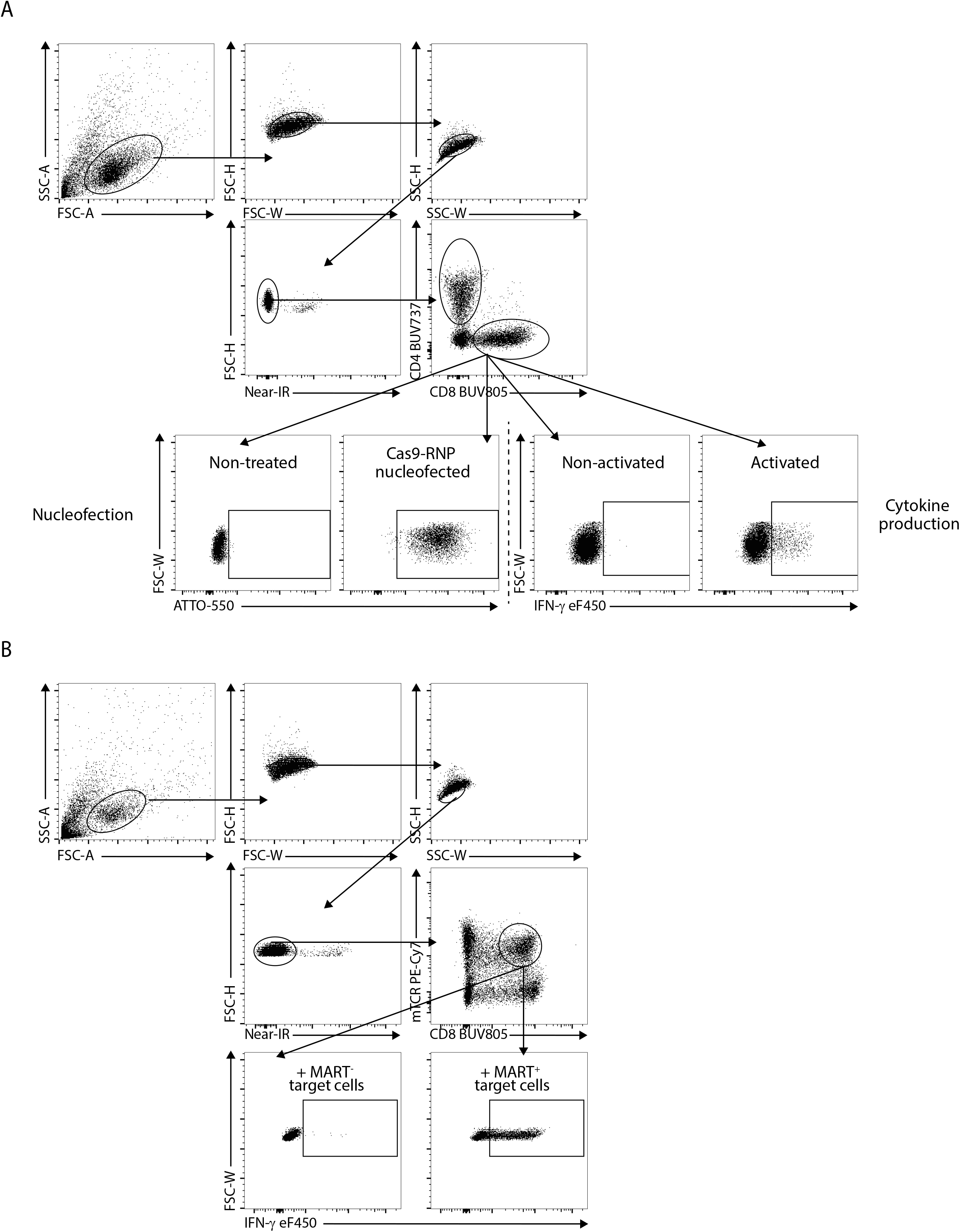
Gating strategies to determine nucleofection and transduction efficiency and T cell cytokine production. Gating strategy for the flow cytometric analysis of human CD8^+^ T cells after nucleofection and antibody-mediated stimulation (A), or after retroviral transduction and coculture with MART-1^+^ and MART-1^−^ target cells (B). In both examples, cells were stained with α-CD4, α-CD8, Near-IR live/dead dye, α-IFN-γ, and in the case of (B) with, α-murine TCR beta. Data were collected with FACSSymphony flow cytometer and analyzed with FlowJo software. Human T cells are identified by their scatter properties (FSC-A x SSC-A plot), and doublets were excluded by gating on FSC-H x FSC-W and SSC-H x SSC-W. Dead cells are excluded based on Near-IR live/dead dye. CD8^+^ T cells are identified by α-CD8 staining. Nucleofected cells are determined based on ATTO550 tracrRNA staining. MART-1 TCR-transduced T cells are determined by α-murine TCR beta staining. In all cases, negative controls or non-stimulated cells were used to set the gates. IFN-γ production is determined by α-IFN-γ staining. TNF-α and IL-2 production are determined in a similar fashion.

**Supplemental Figure 2.**
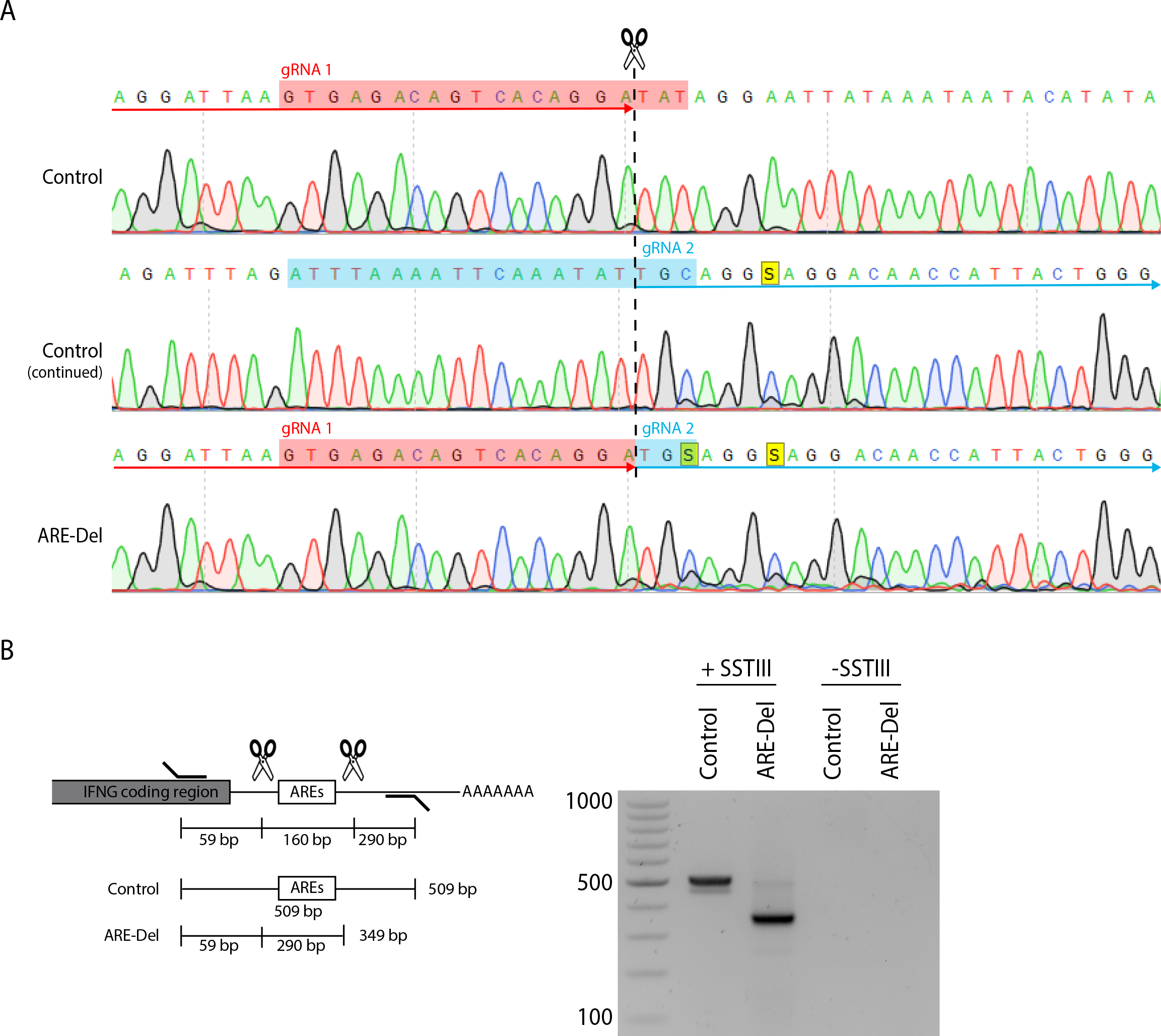
Genomic deletion of the *IFNG* 3’ UTR AREs does not impact 3’ UTR addition. (**A**) Sanger sequencing of control and ARE-Del T cells. crRNA guides are highlighted in red and blue. (**B**) Graphical representation of *IFNG* 3’ UTR in control and ARE-Del T cells, and gel analysis of PCR-amplified *IFNG* 3’UTR from cDNA generated in the presence (+) or absence (−) of SuperScript Reverse Transcriptase (SST) III from DNAse treated nucleotide content from control and ARE-Del T cells.

**Supplemental Figure 3.**
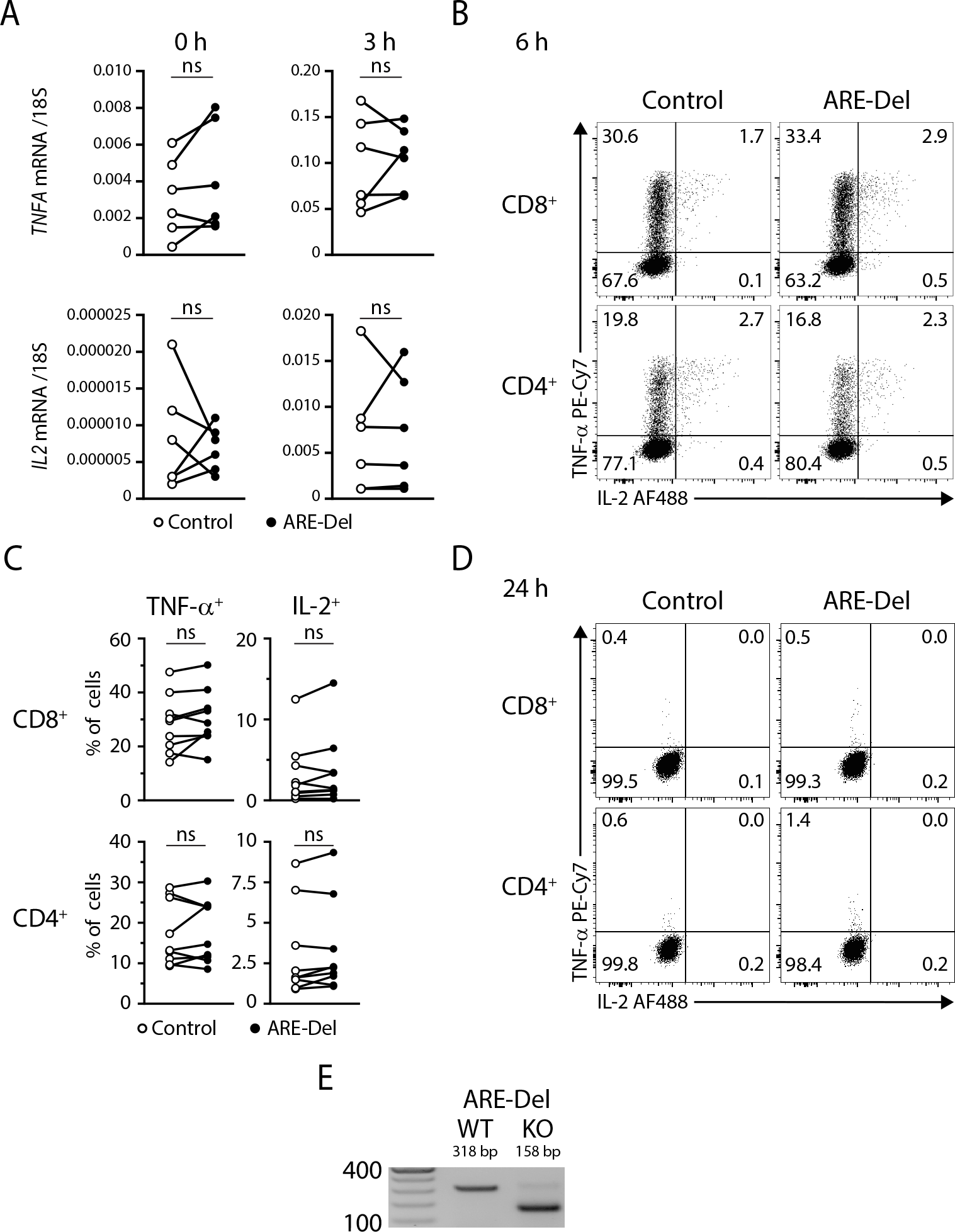
TNF-α and IL-2 mRNA and protein levels are not altered upon *IFNG* ARE-deletion. (**A**) Rested T cells (9 days) were left untreated or restimulated for 3h with α-CD3/α-CD28. *TNFA* and *IL2* mRNA levels from N = 6 donors from 2 independent experiments (paired Student’s t-test; ns, not significant). (**B** and **C**) Resting T cells were restimulated with α-CD3/α-CD28 for 6h. Brefeldin A was added during the last 2h of stimulation. (B) Representative dot plots of TNF-α and IL-2 production of live CD8^+^ and CD4^+^ T cells. (C) Compiled data on TNF-α and IL-2^+^ cells from N = 8 donors from 4 independent experiments (paired Student’s t-test; ns, not significant). T cells cultured in the absence of stimuli were used as negative control. (**D**) Resting T cells were restimulated for 72h with α-CD3/α-CD28 and removed from stimulus for indicated time. TNF-α and IL-2 protein production was assessed by adding Brefeldin A 4h prior to protein production assessment. Representative dot plots of live CD8^+^ and CD4^+^ T cells from n = 7 donors from 3 independent experiments. T cells cultured in the absence of stimuli were used as negative control. (**E**) ARE-deletion in MART-1 TCR-engineered T cells was analyzed by genomic PCR and subsequent gel analysis.

